# Identification and analysis of novel variants in SARS-COV-2 genomes isolated from the Kingdom of Bahrain

**DOI:** 10.1101/2021.01.25.428191

**Authors:** Khalid M bindayna, Abdel Halim Deifalla, Hicham Ezzat Mohammed Mokbel

**Affiliations:** Arabian gulf university; Arabian gulf uninversity

## Abstract

The challenges imposed by the ongoing outbreak of severe acute respiratory syndrome coronavirus-2 affects every aspect of our modern world, ranging from our health to our socio-economic needs. Our existence highly depends on the vaccine’s availability, which demands in-depth research of the available strains and their mutations. In this work, we have analyzed all the available SERS-CoV2 genomes isolated from the Kingdom of Bahrain in terms of their variance and origin analysis. We have predicted various known and unique mutations in the SERS-CoV2 isolated from Bahrain. The complexity of the phylogenetic tree and dot plot representation of the strains mentioned above with other isolates of Asia indicates the versatility and multiple origins of Bahrain’s SERS-CoV2 isolates. We have also identified two high impact spike mutations from these strains which increase the virulence of SARS-CoV2. Our research could have a high impact on vaccine development and distinguishes the source of SERS-CoV2 in the Kingdom of Bahrain.

## Introduction

Severe Acute Respiratory Syndrome CoV-2 or SARS-CoV-2 is the most dangerous threats to humankind across the globe. This disease became a pandemic in early 2020 and effects various countries worldwide. It not only affects the human race by its disastrous impact on human health but also through its devastating influence on every socio-economical aspect of the modern world. It was first reported in China at the city of Wuhan in December 2019. The novel coronavirus, the causative agent of SARS-CoV-2, can be transmitted from man to man through body secretions. SARS-CoV-2 is an enveloped and single-stranded positive-sense RNA containing virus which is closely related to the coronavirus that has been isolated from SARS outbreak in 2003. Though there are some shreds of evidence for the spillover infection of the zoonotic origin of SARS-CoV-2, some controversies also exist regarding its laboratory origin (Andersen et al. 2020).

In the Kingdom of Bahrain, the very first case was detected in 21/02/2020 in a school bus driver who had a travel record to Iran and Dubai. Within three months, total number of cases of SARS-CoV-2 in the Kingdom of Bahrain reaches 10^3^. Though there was a reduction in the increasing rate of new cases after six months, it was again started increasing at the beginning of September. This could be counted as the second wave, which also has started decreasing at the end of November.

This work aims to analyse up to date 150 SARS-CoV-2 genomes that were isolated from the Kingdom of Bahrain. With state of the art comparative genomic approach, we have analysed the mutations present in those strains which will help in the vaccine design. We have also compared all the genomes with twenty different SARS-CoV-2 genomes that were isolated from various parts of Asia, to study the proper origin of SARS-CoV-2 in the Kingdom of Bahrain.

## Method

### Sample selection

We have fetched all the genomes of SERS-CoV2 from the Global Initiative on Sharing All Influenza Data (GISAID) server (Shu et al. 2017). Total 150 genomes with the origin of Bahrain and other 20 genomes with other Asian roots have been identified. The other 20 locations under considerations are Zahedan, Wuhan, Tehran, Semnan, Riyadh, Qom, Qatif, Qatar, Muscat, Makkah, Madinah, Lebanon, Kuwait, Jordan, Jerusalem, Jeddah, Islamabad, Iraq, Dhaka and Delhi. We have only collected complete genome of 28-29MB of sizes.

### Multiple sequence alignment

Multiple sequence alignment has been performed for two groups of the sample. One group contains only the SERS-CoV2 sequences isolated from Bahrain and the other group contains all the genomes from Bahrain and previously mentioned twenty different locations. We performed MSA for both the groups for SNP detection in the first group and phylogenetic tree construction in the second group. Here, we have used MUSCLE (Multiple Sequence Comparison by Log-Expectation) algorithm to align the sequences (Edgar 2004). MUSCLE algorithm is based on two distance measures: k-mer distance and Kimura distance for each of the two sequences. Before multiple alignments, scores for pairwise alignment has been defined. Initially, a binary tree has been created which is followed by accurate multiple alignments in the refinement stage through calculations of time and space complexities.

### Variant and mutations analysis

We have used hCoV-19/Wuhan/WIV04/2019 strain (isolated from Wuhan, China) as reference for variant and mutations analysis of 150 SERS-Cov2 genomes isolated from the Kingdom of Bahrain. We have used CoVsurver mutant analysis tool from GISAID server. This tool identifies specific mutations of SERS-Cov2 along with various structural and functional proteins. Similarly, for characterization of Single Nucleotide Polymorphisms (SNPs), we have used SNiPlay tools available at (http://sniplay.cirad.fr/.). This tool helps us to in the determination of Single Nucleotide Polymorphisms (SNPs) and insertion/deletion (indels) along the whole genome length. Both the tools accept aligned FASTA sequence.

### Phylogenetic tree construction

Phylogenetic tree construction is required for the detection of similarities in SERS-Cov2 genomes isolated from various places. it is also helpful to detect the close relative or source of the 150 different SERS-Cov2 genomes isolated from the Kingdom of Bahrain. For phylogenetic tree reconstruction, we have used the PhyML algorithm which is based on the maximum likelihood method (Guindon 2003). It allows very large datasets in multiple sequence aligned format. This method follows the hill-climbing algorithm which simultaneously adjusts tree topology and branch lengths. It starts with a basic tree buildup via the fast distance-based method and with every iteration, it improves the likelihood of the tree. It is a very fast method which reaches optima within a few iterations.

### Dot plot assay

The Dot plots are used to compare large sequence sets to get an insight into the similarity overview. A distinct set of sequences will produce a diagonal straight line. We have used D-genies (Cabanettes 2018) which is under GNU General Public License (GPL). Here, similar sequences will produce a scatter plot as most of the sequences will be matched with each other. A large number of samples could be plotted with a multiple sequence aligned format as an input file. We have used this plot to estimate the overall 170 previously mentioned SERS-Cov2 sequences to check their similarities and relatedness.

## Results

### Mutation analysis

We have compared total 150 genomes that have been isolated from the Kingdom of Bahrain with the reference genome from Wuhan strain (hCoV-19/Wuhan/WIV04/2019) which is the official reference sequence employed by GISAID. We have analysed all the mutations in the 150 genomes compared to the reference strain. A detailed list of unique and known mutations is given in Supplementary Information 1. The unique mutations are not reported previously. Here, we have listed down eight strains which have >= 5 unique mutations in their genomes in Table 1.

**Table 1:**
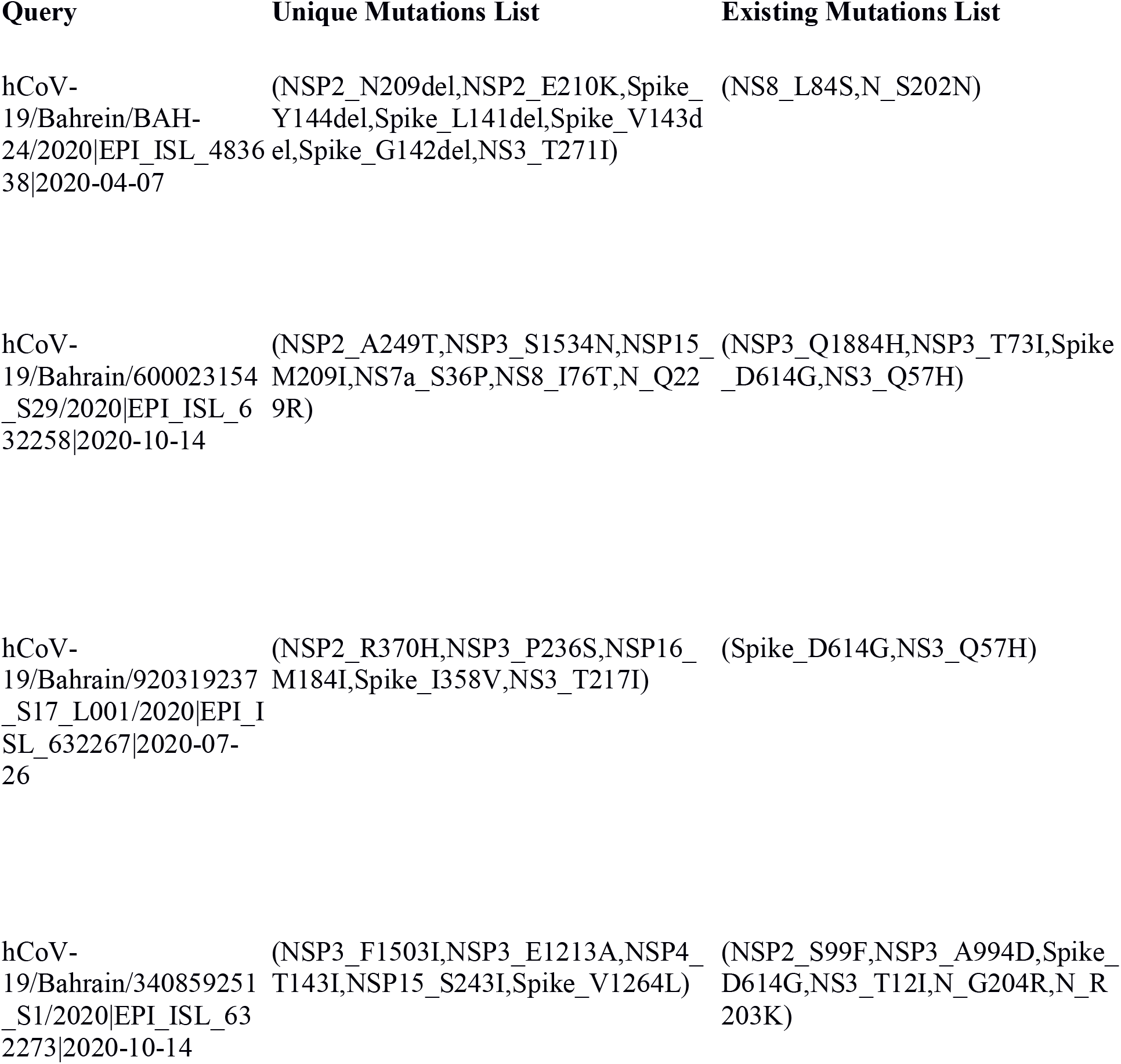
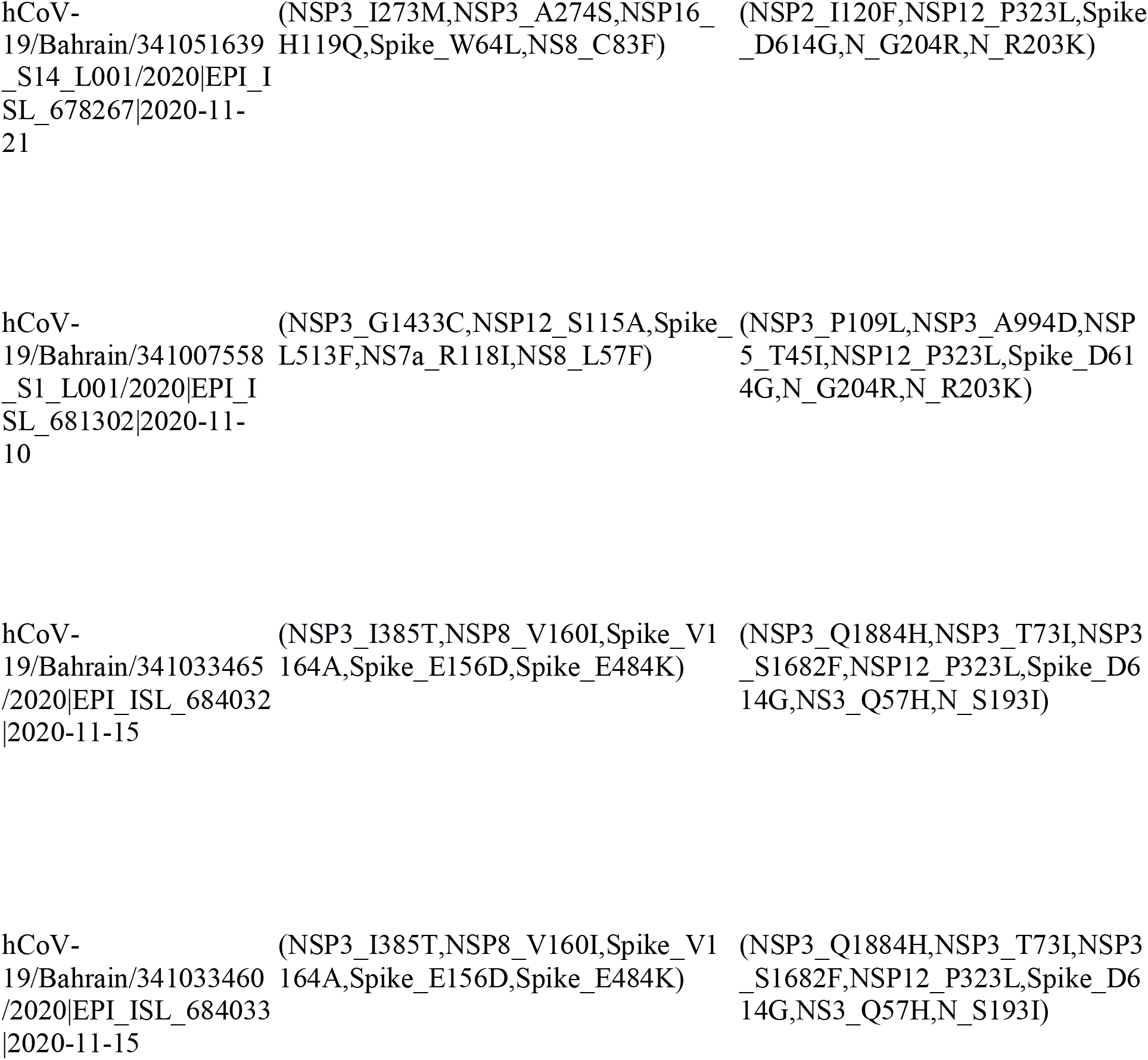
Eight strains isolated from the kingdom of Bahrain which have >= 5 unique mutations in their genomes.

We have also summarized the positions of these mutations on the spike glycoprotein in the unbound state (PDB id: 6ACC) and bound state (PDB id: 6ACJ) with host ACE2 receptor (green) in Fig 1 (A) and (B) respectively (Ortega et al. 2020, Zhang et al. 2020).

**Fig 1.**
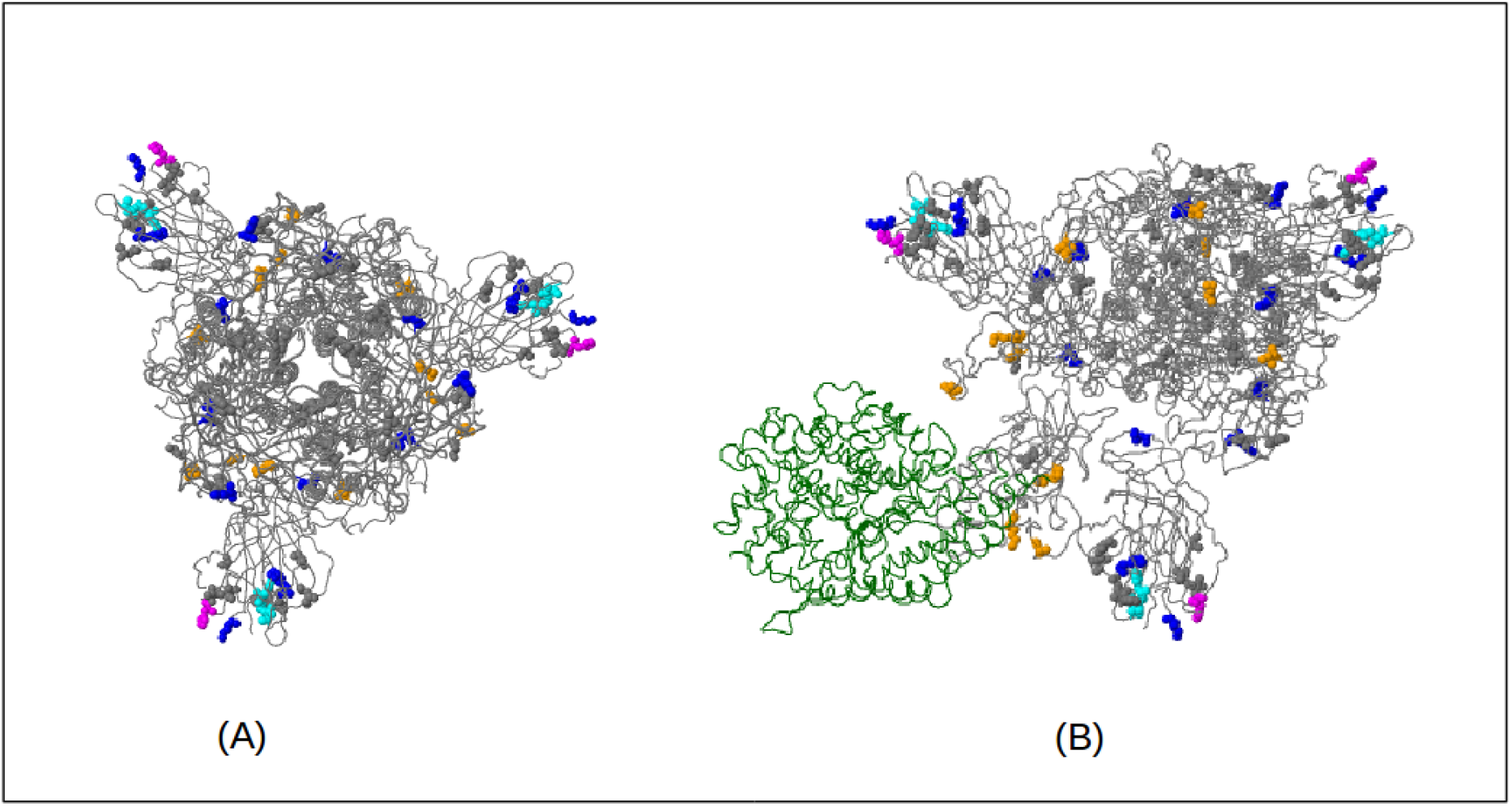
Mutations on the spike glycoprotein in the unbound state (PDB id: 6ACC) and bound state (PDB id: 6acj) with host ACE2 receptor (green) in Fig 1 (A) and (B) respectively.

Mutations on the spike glycoprotein that were previously identified have been colour coded in the given structure. H49Y[blue], W64L, A67S, G75S(77), T76I(77)[magenta], R78M, R102I[blue], V127F, L141del[sky], G142del[sky], V143del[sky], Y144del(143)[sky], E154G, E156D, M177I, S255F(257)[blue], I358V[yellow], L452M[yellow], S459F[yellow], E484Q[yellow], E484K[yellow], L513F, D614G[blue], V622I, Q675H(674)[blue], S689I(692), A871S, I909V, L1063F, E1092A and H1101Y. Stain hCoV-19/Bahrein/BAH-24/2020|EPI_ISL_483638|2020-04-07 is identified to posses maximum number of unique mutations. The analyzed the genome variations in terms of SNP’s for these 150 genomes are given in Supplementary Information 2. Among these mutations, E484K and D614G are recently noted as the significant mutations due to their effect on SARS-CoV 2 virulence. E484K mutation on the spike protein causes SARS-CoV 2 escape from the neutralizing effect of the human immune systems mediated by antibody (Mahase 2021). On the other hand, D614G mutation on the spike protein has high infectivity in human cells having ACE2 receptors (Ogawa et al. 2020, Kim et al. 2020). D614G mutation situated exactly at the receptor binding domain of the spike protein. So due to this mutation, affinity of ACE2 receptor in increases. It was reported that infectivity increases by >1/2 log_10_ which is measured through cell fusion assay (Ogawa et al. 2020).

### Phylogenetic tree analysis

We have reconstructed phylogenetic tree separately for the SERS-Cov2 sequences isolated from the Kingdom of Bahrain and all 170 sequences from all over Asia. Here, first of all our main aim was to identify the similarities between SERS-Cov2 sequences isolated from the Kingdom of Bahrain. A view of phylogenetic tree only for Bahrain sequences has been presented in Fig 2. It can be observed from Fig 2 that there is a huge diversity in the genomic sequences isolated from various places of Bahrain. This could definitely increases our curiosity for finding the actual origin of SERS-Cov2 in Bahrain. So we constructed the second tree which posses all the 170 sequences from Bahrain as well as the sequences from various places of Asia. The phylogenetic tree for all the 170 sequences is represented in Fig 3. From Fig 3. it seems that the origin of SERS-Cov2 in Bahrain is not only from one place. There is huge diversity in the phylogenetic tree where we can see that 37 sequence of SERS-Cov2 isolated from Bahrain is very much different from other sequences. Other 113 sequences have different origin. Some of the sequence have similarities with Qatif where some other sequences have similarities with the sequence isolated from Iran, Qatar and Israel. Such huge diversity leads us to perform a dot plot assay for overall understanding of all the 170 sequences.

### Dot plot assay

We have placed all 170 sequences in a specific manner both in the X and the Y axis of a graph, and study the similarity matrix between them. The dot plot is represented in Fig 4.

**Fig 4.**
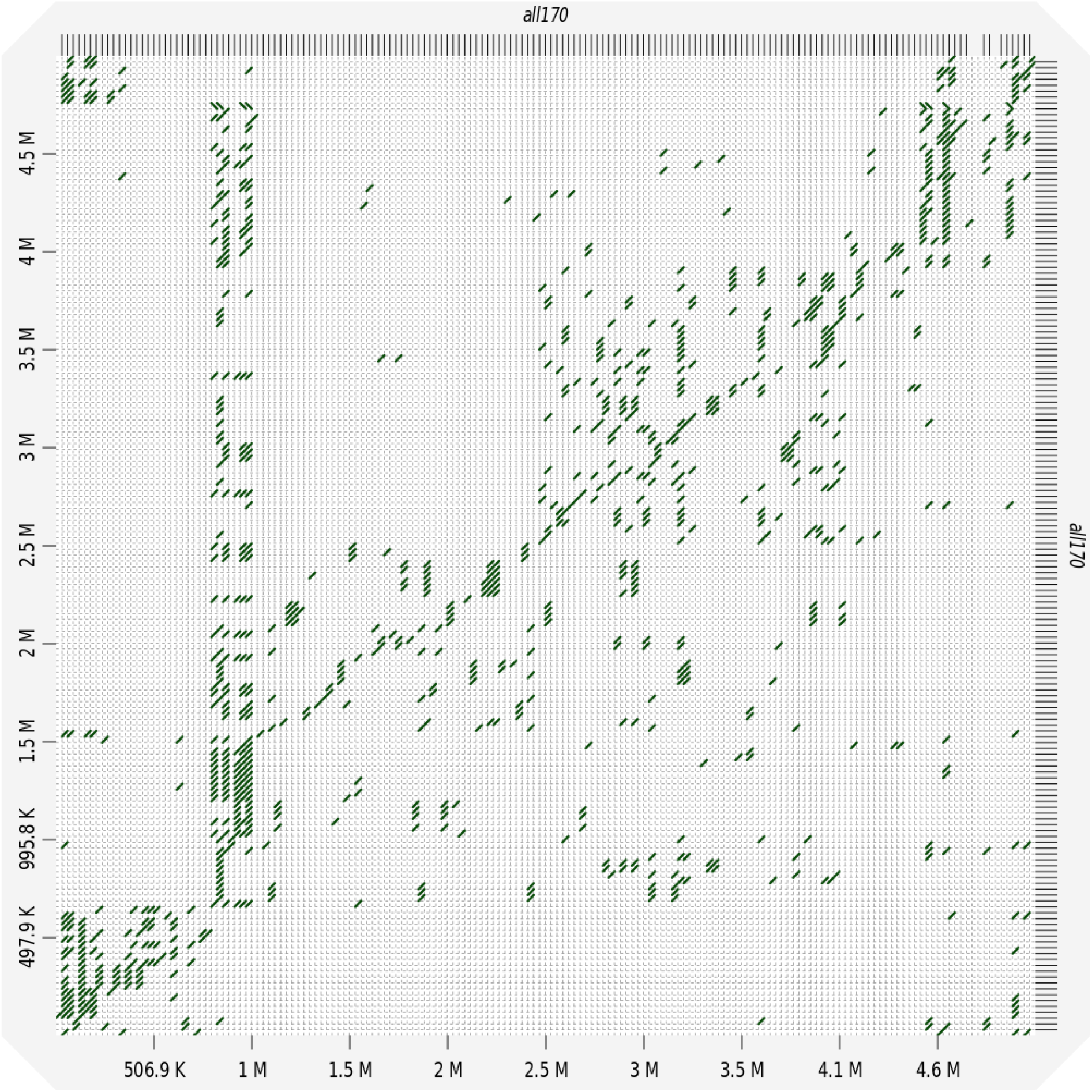
A dot plot assay of all the 170 sequences isolated from various regions of Asia.

The scattered nature of the dot plot in Fig 4 clearly indicates that all the 170 genomes are closely related to each other. If there is a single origin of the 150 SERS-Cov2 genomes isolated from Bahrain, then the plot looks like a diagonal straight line. So we can conclude that the origin of SERS-Cov2 in Bahrain is not only from one place.

## Discussion

In this paper we have represented a detailed account of total 150 SERS-Cov2 genomes that have been isolated from the Kingdom of Bahrain. We have analyzed the mutations in all the strains in reference with the hCoV-19/Wuhan/WIV04/2019 Wuhan strain which is considered as the origin or SERS-Cov2. We have identified various known and unknown mutations from these 150 strains. We saw that hCoV-19/Bahrein/BAH-24/2020|EPI_ISL_483638|2020-04-07 possesses maximum number of unique mutations. We have found two significant mutations (E484K and D614G) on the spike protein of Bahrain isolates. These mutations cause decrease in antibody affinity towards SARS-CoV2 and increase of binding affinity towards ACE2 receptor respectively. A detailed account of all these mutations will be helpful for designing the vaccine against SERS-Cov2. We have also studied the origin of SERS-Cov2 in the Kingdom of Bahrain. We found that there is a huge diversity in the 150 genomes. This indicates that there could be multiple source of SERS-Cov2 in the Kingdom of Bahrain.

## References

(1) Andersen, K. G., Rambaut, A., Lipkin, W. I., Holmes, E. C., & Garry, R. F. (2020). The proximal origin of SARS-CoV-2. Nature medicine, 26(4), 450–452. https://doi.org/10.1038/s41591-020-0820-9

(2) Shu, Y., & McCauley, J. (2017). GISAID: Global initiative on sharing all influenza data–from vision to reality. Eurosurveillance, 22(13), 30494.

(3) Edgar, R. C. (2004). MUSCLE: multiple sequence alignment with high accuracy and high throughput. Nucleic acids research, 32(5), 1792–1797.

(4) Guindon S, Gascuel O. A simple, fast, and accurate algorithm to estimate large phylogenies by maximum likelihood. Syst Biol. 2003 Oct;52(5):696–704. doi: 10.1080/10635150390235520. PMID: 14530136.

(5) Cabanettes F, Klopp C. (2018) D-GENIES: dot plot large genomes in an interactive, efficient and simple way. PeerJ 6: e4958

(6) Ortega, J. T., Serrano, M. L., Pujol, F. H., & Rangel, H. R. (2020). Role of changes in SARS-CoV-2 spike protein in the interaction with the human ACE2 receptor: An in silico analysis. EXCLI journal, 19, 410.

(7) Zhang, C., Zheng, W., Huang, X., Bell, E. W., Zhou, X., & Zhang, Y. (2020). Protein structure and sequence reanalysis of 2019-nCoV genome refutes snakes as its intermediate host and the unique similarity between its spike protein insertions and HIV-1. Journal of proteome research, 19(4), 1351–1360.

(8) Mahase, E. (2021). Covid-19: What new variants are emerging and how are they being investigated?.

(9) Ogawa, J., Zhu, W., Tonnu, N., Singer, O., Hunter, T., Ryan, A. L., & Pao, G. M. (2020). The D614G mutation in the SARS-CoV2 Spike protein increases infectivity in an ACE2 receptor dependent manner. Biorxiv.

(10) Kim, S., Lee, J. H., Lee, S., Shim, S., Nguyen, T. T., Hwang, J.…, & Kim, S. (2020). The progression of sars coronavirus 2 (sars-cov2): Mutation in the receptor binding domain of spike gene. Immune Network, 20(5).

